# Attack behaviour in naïve Gyrfalcons is modelled by the same guidance law as in Peregrines, but at a lower guidance gain

**DOI:** 10.1101/2020.10.06.328260

**Authors:** Caroline H. Brighton, Katherine E. Chapman, Nicholas C. Fox, Graham K. Taylor

**Author notes:** **CORRESPONIDNG AUTHORS:**.

## Abstract

The aerial hunting behaviours of birds are strongly influenced by their flight morphology and ecology, but little is known of how this variation relates to the behavioural algorithms guiding flight. Here we use onboard GPS loggers to record the attack trajectories of captive-bred Gyrfalcons (*Falco rusticolus*) during their maiden flights against robotic aerial targets, which we compare to existing flight data from Peregrines (*Falco peregrinus*) The attack trajectories of both species are modelled most economically by a proportional navigation guidance law, which commands turning in proportion to the angular rate of the line-of-sight to target, at a guidance gain *N*. However, Gyrfalcons operate at significantly lower values of *N* than Peregrines, producing slower turning and a longer path to intercept. Gyrfalcons are less agile and less manoeuvrable than Peregrines, but this physical constraint is insufficient to explain their lower guidance gain. On the other hand, lower values of *N* promote the tail-chasing behaviour that is typical of wild Gyrfalcons, and which apparently serves to tire their prey in a prolonged high-speed pursuit. Moreover, during close pursuit of fast evasive prey such as Ptarmigan (*Lagopus* spp.), proportional navigation will be less prone to being thrown off by erratic target manoeuvres if *N* is low. The fact that low-gain proportional navigation successfully models the maiden attack flights of Gyrfalcons suggests that this behavioural algorithm is embedded in a hardwired guidance loop, which we hypothesise is ancestral to the clade containing Gyrfalcons and Peregrines.

**SUMMARY STATEMENT:** Naïve Gyrfalcons attacking aerial targets are modelled by the same proportional navigation guidance law as Peregrines, but with a lower navigation constant that promotes tail-chasing rather than efficient interception.

## INTRODUCTION

Raptorial feeding is a complex mode of foraging behaviour, the success of which hinges on intercepting a target whose own success hinges on evading capture. Aerial pursuit in particular is one of the most challenging behaviours that organisms perform, but also one of the simplest to characterise. A dyadic interaction, for example, is minimally described by the trajectories of a pair of interacting particles representing the predator and its prey. This level of description lends itself to an algorithmic approach, in which a mathematical rule – in this case, a particular kind of behavioural algorithm known as a guidance law – is used to connect sensory input to motor output (Hein et al. 2020). As there are only a limited number of ways in which one particle can be steered to intercept another, this algorithmic approach lends itself in turn to a rigorous comparative analysis of behaviour across different taxa, locomotor modes, and spatiotemporal scales. Key research questions include: What sensory information is used to guide interception, and how? For what function is the attacker’s guidance algorithm optimized? And how is this behavioural algorithm acquired? Here we address these questions for a sample of naïve Gyrfalcons (*Falco rusticolus*), which are the largest of all falcons, and hence one of the largest extant predators specialising in aerial interception.

Gyrfalcons are closely related to Peregrines (*Falco peregrinus*), whose attack trajectories are well modelled (Brighton et al. 2017) by a guidance law called proportional navigation (PN). A pure PN guidance law commands turning at an angular rate 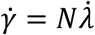, where 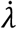 is the angular rate of the attacker’s line-of-sight to target, and where the guidance gain *N* is called the navigation constant and assumed to be fixed within an attack. In contrast, the attack trajectories of Harris’ Hawks (*Parabuteo unicinctus*) are best modelled by a mixed PN+PP guidance law, 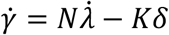, which combines a PN element with a proportional pursuit (PP) element, −*Kδ*, commanding turning in proportion to the deviation angle *δ* between the attacker’s velocity vector and its line-of-sight to target (Brighton and Taylor 2019). In this case, a PN+PP guidance law with guidance constants *N* = 0.7 and *K* = 1.2 s^−1^ modelled the observed flights more closely than a PN or PP guidance law in which *N* or *K* was allowed to vary between flights (Brighton and Taylor 2019). The PN guidance law of Peregrines promotes short-cutting towards the eventual point of intercept, making it well suited to intercepting non-manoeuvring targets in open environments (Brighton et al. 2017). In contrast, the PN+PP guidance law of Harris’ Hawks promotes tail-chasing directly after the target, making it better suited to close pursuit through potentially cluttered environments (Brighton and Taylor 2019). Hawks (Accipitridae) and falcons (Falconidae) are thought to have diverged >60 mya (Jarvis et al. 2014; Prum et al. 2015), so we would naturally expect the attack behaviour of Gyrfalcons to be better modelled by the PN guidance law of Peregrines than by the PN+PP guidance law of Harris’ Hawks.

Peregrines and Gyrfalcons are both adapted to open environments, hunting mainly avian prey, which may be knocked down in flight (Garber et al. 1993), struck on the ground before taking flight (Bengtson 1971), or forced to the ground after a long chase (Cade 1982; Woodin 1967). However, whereas Peregrines will often dive from altitude in a high-speed stoop (Cresswell 1996), Gyrfalcons rarely stoop in the wild and almost always hunt close to the ground (Cade 1982; Garber et al. 1993). Gyrfalcons are most often recorded performing low surprise attacks initiated from a perch or from ridge soaring (Cade 1982; Garber et al. 1993; Potapov and Sale 2005; White 1991; White and Weeden 1966), but if not immediately successful then they will commonly enter the prolonged tail-chase that is typical of this species (Cade 1982; Pennycuick et al. 1994). Similar hunting behaviours are also observed in Peregrines (Cresswell 1996), but the distinct speed advantage that Peregrines acquire when stooping (Mills et al. 2018; Mills et al. 2019) appears to be lacking in wild Gyrfalcons. Such variation in hunting behaviour may be expected to be associated with variation in the underlying guidance law.

Peregrine attacks are best modelled by a range of values of *N* (median: 2.6; 1^st^, 3^rd^ quartiles: 1.5, 3.2) lower than the interval 3 ≤ *N* ≤ 5 that is typical of missile applications, but close to the optimum minimizing total steering effort in the classical linear-quadratic formulation of the optimal guidance problem (Brighton et al. 2017). This theory predicts that *N* = 3 *v_c_*/(*v* cos *δ*) is optimal for attacks on non-manoeuvring targets, where *v_c_* is the speed at which the attacker closes range on its target, and *v* is the attacker’s groundspeed (Shneydor 1998; Siouris 2004). In words, the optimal value of *N* is proportional to the ratio *v_c_*/(*v* cos *δ*), which expresses how effectively the attacker closes range on its target (*v_c_*, representing the difference between the speed of the attacker’s approach and the speed of the target’s retreat) in relation to its own motion towards the target (*v* cos *δ*, representing the speed of the attacker’s approach). Hence, whereas *N* = 3 is optimal for attacks on stationary targets (where *v_c_ = v* cos *δ*), *N* < 3 is optimal for attacks on retreating targets (where *v_c_ < v* cos *δ*), at a value which depends on the relative speeds of target and attacker. In a high-speed stoop, for instance, the target’s speed may be negligible compared to that of its attacker, such that *v_c_*/(*v* cos *δ*) ≈ 1 making *N* ≈ 3 optimal (Mills et al. 2018; Mills et al. 2019). Conversely, in a prolonged tail chase in which the target flees at a similar speed to the attacker, much lower values of *N* < 3 may be optimal. We therefore hypothesise that Gyrfalcons will use PN guidance to intercept targets but will do so using a lower value of *N* than Peregrines.

## METHODS

To test these hypotheses, we use a combination of empirical measurements and computational modelling to identify the guidance law used by naïve captive-bred Gyrfalcons chasing robotic aerial targets. Our observations were recorded on the first attack flights that the birds had ever made against aerial targets, and therefore reflect as closely as possible the hard-wired form of the underlying guidance algorithm, with no prior opportunity for reinforcement learning.

### Animals

We observed 23 naïve captive-bred Gyrfalcons, comprising 19 pure Gyrs (*F. rusticolus*) and 4 Gyr-Saker hybrids (7/8^th^ *F. rusticolus* × 1/8^th^ *F. cherrug*), chasing robotic aerial targets during their first flight sessions. This sample contained only naïve first-year birds that had not previously flown after aerial targets, except for during a single flight against a swung lure immediately beforehand. This work has received approval from the Animal Welfare and Ethical Review Board of the Department of Zoology, University of Oxford in accordance with University policy on the use of protected animals for scientific research, permit no. APA/1/5/ZOO/NASPA, and is considered not to pose any significant risk of causing pain, suffering, damage or lasting harm to the animals.

### Experimental protocol

Flight trials were carried out on open moorland at Watch Hill, Wealside Farm, Northumberland, UK, in winds gusting from 4 to 7 m·s^−1^. The birds were recorded chasing a remotely piloted, ducted fan “Roprey” model with a food reward strapped to its dorsal surface (Wingbeat Ltd, Carmarthen, Wales, UK; Fig. 1). Each bird carried a BT-Q1300 GPS receiver (QStarz International, Taipei, Taiwan) logging position and groundspeed at 5 Hz. The GPS receiver was carried dorsally on a Trackpack harness (Marshall Radio Telemetry, Salt Lake City, UT, USA), giving a total load of 0.015 kg. An identical GPS logger was fixed inside the body compartment of the Roprey, and the flights were filmed using a handheld Lumix DMC-FZ1000 camera recording 4k video at 25 frames s^−1^ (Panasonic Corporation, Osaka, Japan). Each flight trial began as the falconer unhooded the bird on their fist, with the Roprey held ~20 m upwind. The Roprey was launched as soon as the bird took off, or sometimes just beforehand; if not caught immediately (see e.g. Fig. 2A), it was flown through a series of evasive turns (see e.g. Fig. 2B). The trial ended when the bird first intercepted the Roprey. If the bird knocked the Roprey with its talons, then the pilot brought it safely to the ground; if the bird bound to the Roprey, then the motor was cut, and an airbrake was deployed to prevent the pair from drifting. In a few cases, the Roprey crash-landed before finally being intercepted by the bird. After disregarding 3 flights in which the bird did not intercept the target, and another 7 flights in which the GPS logger was lost during the session, we were left with an initial sample of 28 flights from 19 naïve Gyrfalcons.

**Figure 1.**
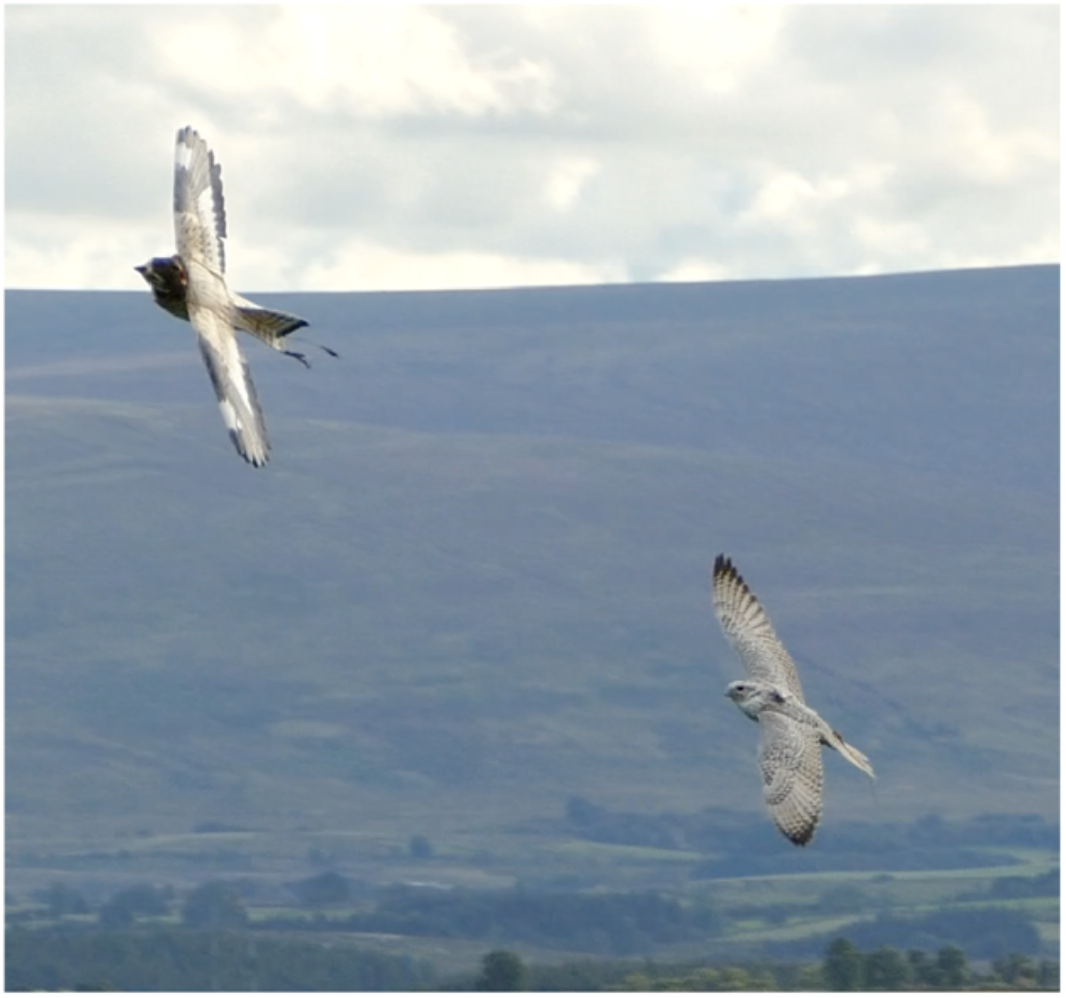
Cropped frame from video of a typical chase involving a Gyrfalcon and a “Rokarrowan” Roprey model. Note the proximity of the attacker to its target, and their similar bank angles, which are characteristic of the tail-chasing behaviour that we observed.

**Figure 2.**
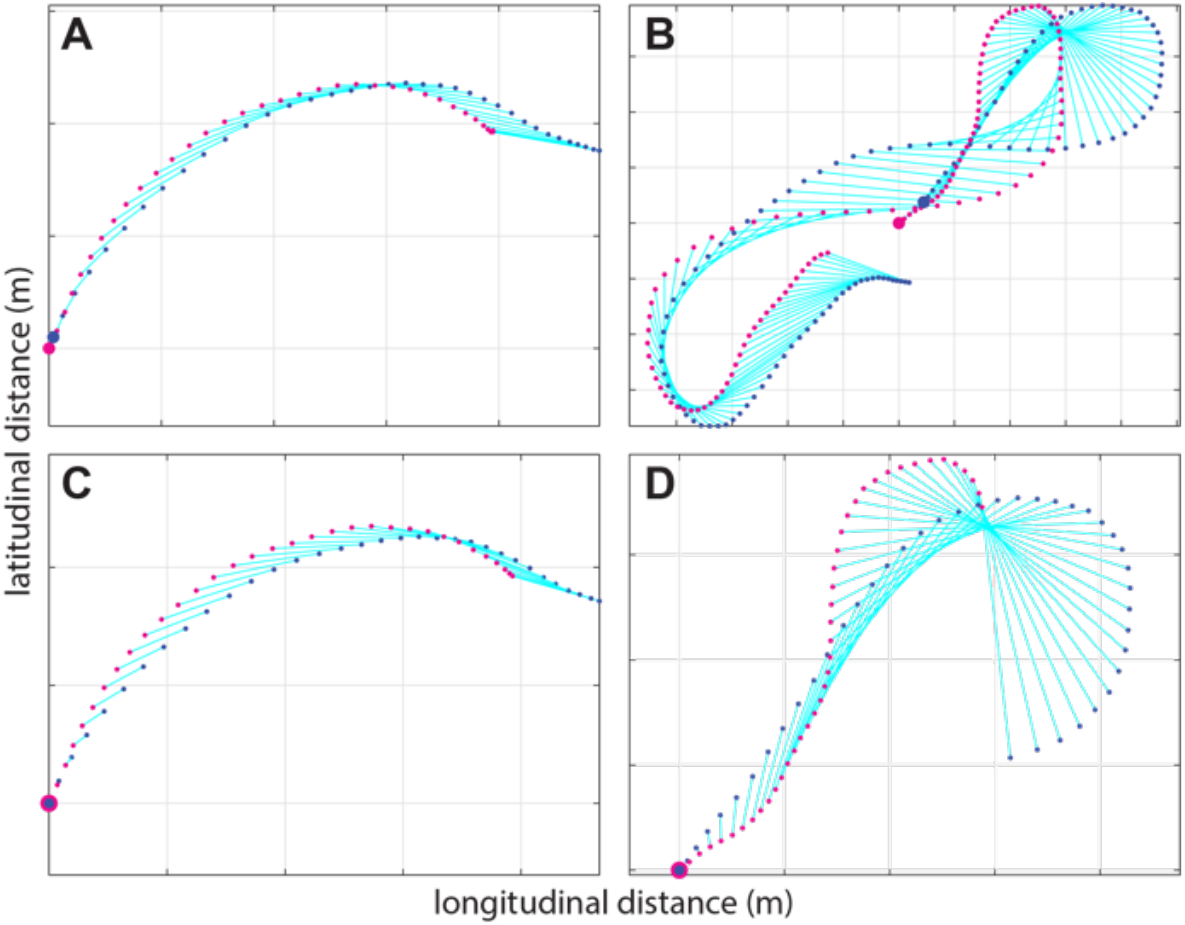
Two-dimensional (2D) GPS trajectories for: (A) the entirety of a short dash; and (B) the entirety of an extended chase, showing the lines-of-sight (cyan lines) between the Gyrfalcon (blue points) and Roprey (magenta points) at each sample point. Note the small discrepancy in the estimated position of target and attacker at the point of intercept (enlarged sample points), expected because of the positioning error associated with GPS receivers (see Methods). (C,D) The terminal phase of the same two flights (see Fig. 4G,M for modelling), after shifting the attacker’s trajectory to correct for this positioning error. Gridlines at 10m spacing.

### Data synchronization and error analysis

We used the GPS time signal to synchronize the two GPS data streams to within ± 0.1 s, having linearly interpolated a small number of dropped datapoints. We matched the synchronized GPS data to the video with reference to take-off and landing, and used the video to identify the time of first intercept. We then shifted the entire GPS trajectory of the bird so as to match its estimated position at first intercept to that of the target (Fig. 2). This adjustment was necessary to remove positional bias due to GPS receiver clock error, which is such that at the nominal horizontal positioning accuracy of our GPS receivers (<3.0 m circular error probable), we would expect 50% of position estimates of a co-located pair of receivers to be separated by ≥4.8 m under an isotropic gaussian error model. Receiver clock error varies slowly once settled, so has no significant effect on the measurement of changes in position over short timescales: in fact, we have shown empirically that our receivers have a precision of order 0.1 m for changes in position occurring over intervals of order 10 s (Brighton et al. 2017). Even so, there were 8 flights for which the discrepancy in the horizontal position estimates of the bird and target at intercept exceeded the 95^th^ percentile expected at the nominal accuracy of the receivers, which indicates a higher-than-expected positioning error in one or both receivers. Of these 8 flagged flights, 5 were the first flight that we recorded after receiver start-up at the beginning of a logging session, which suggests that the receiver clock estimate had not been given sufficient time to settle at the start of all 7 logging sessions (Fisher’s exact test: *p* = 0.01). We therefore dropped these 8 flagged flights from the analysis, leaving a final sample of 20 flights by *n* = 13 Gyrfalcons (Table S1). Among these 20 remaining flights, the discrepancies in the position estimates of bird and target at intercept (median: 5.5 m; IQR: 8.0-3.1 m) were distributed as expected at the nominal accuracy of the receivers (median: 4.8m; IQR: 7.3-2.8 m), albeit with no extreme outliers.

### Trajectory modelling

We simulated the birds’ measured flight trajectories in Matlab, by predicting their turn rate 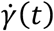 in response to the target’s measured trajectory using a guidance law of the form:

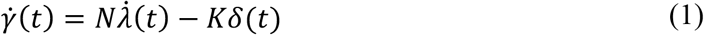

where 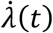 is the angular rate of the line-of-sight to target, where *δ*(*t*) is the deviation angle between the target and the attacker’s velocity vector, and where *N* and *K* are constants. Angles and angular rates are defined in vector form in the three-dimensional (3D) case, but can be treated as scalars in the two-dimensional (2D) case. Eq. 1 necessarily ignores the effects of any sensorimotor delay, on the basis that this is expected to be comparable to the ±0.1 s uncertainty in the synchronization of the GPS data streams; see (Brighton and Taylor 2019). In the special case that *K* = 0, Eq. 1 reduces to a pure PN guidance law, whereas in the special case that *N* = 0, Eq. 1 reduces to a pure PP guidance law. We refer to the general case in which *K* ≠ 0 and *N* ≠ 0 as mixed PN+PP guidance. We used a forward Euler method to simulate each flight under either PN, PP, or PN+PP guidance, initializing the simulation using the bird’s measured position and velocity at some given start point, and matching the bird’s simulated flight speed to its known groundspeed. The method is described further in our work with Peregrines and Harris’ Hawks (Shneydor 1998; Siouris 2004), the only difference being the 1/3000^th^ s step size that was used here to ensure that all parameter estimates were accurate at the reported precision (for simulation code, see Supporting Data S1).

We used a Nelder-Mead simplex algorithm to find the value of *N* and/or *K* that minimized the mean prediction error for each flight, defined as the mean absolute distance between the measured and simulated trajectories. However, because the bird did not always start chasing the target from the moment it was launched, it was necessary to select the start-point of the simulation by reference to the data. Following the same approach used to model aerial chases in Peregrines (Brighton et al. 2017), we therefore ran simulations beginning from all possible start times ≥2.0 s before intercept, reporting the longest 2D simulation (up to a maximum of 20 s) for which the mean prediction error was ≤1.0% of the flight distance modelled. For the 3D case, we used an equivalent error tolerance of 1.2%, to preserve the same tolerance in each dimension. This process of selecting the start-point of each simulation by reference to the model’s performance on the data is objective, but risks capitalizing on chance. We ensure that our inferences are robust to the associated risk of overfitting by making contrastive inferences between alternative guidance models and by focussing our inferences on the population properties of the estimated parameters. Simulations that failed to model ≥2.0 s of flight at the specified error tolerance are recorded as unsuccessful, and their parameter estimates are not reported. Hence, because all of the successful simulations were fitted to within the same specified error tolerance, the appropriate figure of merit for each simulation is the overall distance or duration of flight modelled, expressed relative to the number of estimated guidance parameters.

To facilitate direct comparison with our published results from experienced Peregrines (Brighton et al. 2017), we reanalysed the Peregrine dataset using exactly the same methods described above. After excluding 33 flights representing attacks on stationary ground targets, and after dropping another 9 flights that were flagged as having lower-than-expected accuracy at the point of intercept by the method above, this yielded a refined sub-sample of 13 flights against aerial targets made by 4 experienced Peregrines (*F. peregrinus*). The mixed PN+PP guidance law was not considered in the original analysis (Brighton et al. 2017), so was fitted for the first time here. The only other difference from the original analysis was the refinement of the integration step size used here. This had only a small effect on the simulated trajectories, but sometimes caused a different start-point to be selected for the simulations, because of the thresholding associated with finding the longest simulation fitted at the specified error tolerance.

## RESULTS

### The attack trajectories of naïve Gyrfalcons are well modelled under PN

The 2D simulations under PN successfully modelled 18/20 of the flights by naïve Gyrfalcons, fitting 1127 m and 111.0 s of flight at 1.0% error tolerance (Figs. 2A-B, 3, S1; Table S2). In contrast, the PP simulations modelled only 14/20 flights successfully, fitting 824 m and 77.4 s of flight, for the same number of estimated guidance parameters (Fig. 3A-B; Table S2). PN is therefore much the better supported of the two pure guidance laws. In contrast, the PN+PP simulations successfully modelled the terminal phase of all 20 flights, fitting 1358 m and 139.4 s of flight at 1.0% error tolerance (Fig. 3A-B; Table S2). The addition of a PP term therefore increased the duration of flight fitted by a factor of 1.3 relative to PN, for a doubling in the number of estimated guidance parameters. It follows that the Gyrfalcons’ attack trajectories are more economically modelled by PN than by PN+PP.

**Figure 3.**
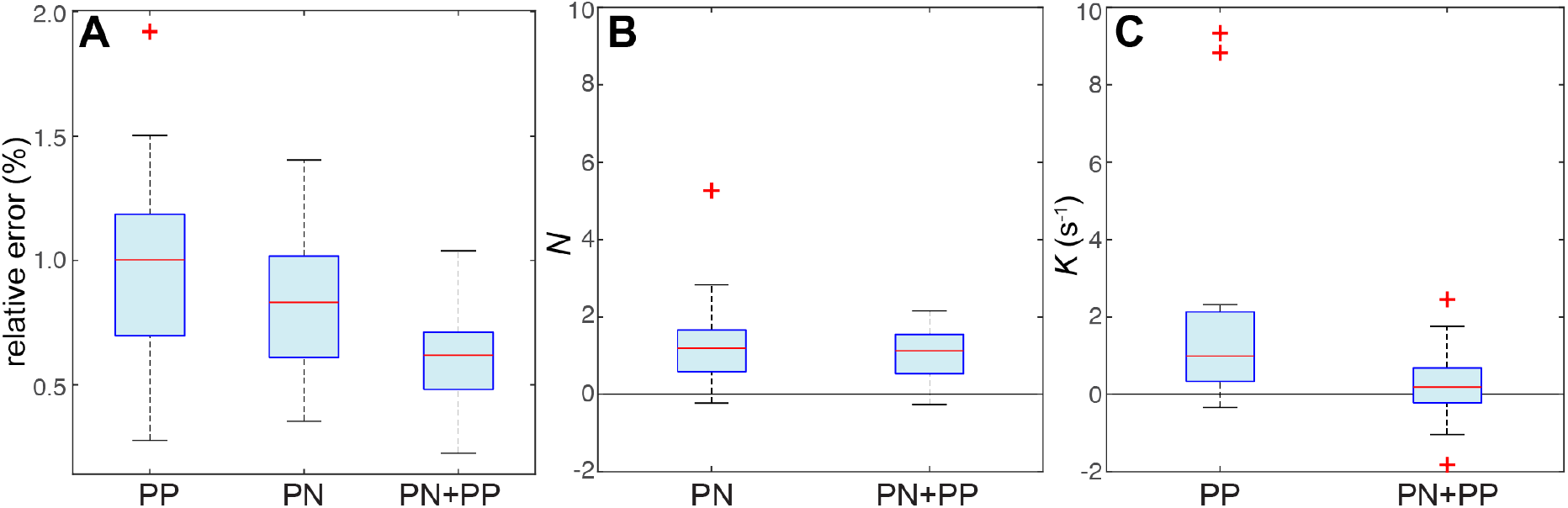
Box-and-whisker plots comparing model fits and parameter estimates for the 2D simulations of 20 flights from *n* = 13 Gyrfalcons under proportional pursuit (PP), proportional navigation (PN), and mixed (PN+PP) guidance. (A) Relative error of simulation, showing either the relative error for the longest simulation lasting ≥ 2 s that met the 1.0% error tolerance threshold for each flight, or if no simulation met this threshold, then the minimum relative error achieved on any simulation lasting ≥ 2 s. Note that whilst the PN+PP simulations fit the flights more closely than either PN or PP, they are almost certainly overfitted (see Results). (B,C) Parameter estimates for the guidance constants *N* and *K* for all successfully modelled flights. Note the variable sign of the parameter estimates for *K* under PN+PP, which confirms that these simulations are overfitted (one outlier for PN+PP not shown). The centre line of each box denotes the median for all flights; the lower and upper bounds of the box denote the 1^st^ and 3^rd^ quartiles; crosses indicate outliers falling >1.5 times the interquartile range beyond the 1^st^ or 3^rd^ quartile; whiskers extend to the farthest datapoints not treated as outliers.

The implied overfitting of the PN+PP simulations is further evidenced by the observation that the parameter estimates for *K* were inconsistently signed in the PN+PP simulations (median *K*: 0.2 s^−1^; 1^st^, 3^rd^, quartiles: −0.2, 0.7 s^−1^; sign test: *p* = 0.26; *n* = 20 flights; Fig. 3D), despite being positive in most of the successful PP simulations (median *K*: 0.9 s^−1^; 1^st^, 3^rd^ quartiles: 0.2, 1.7 s^−1^; sign test: *p* = 0.01; *n* = 14 flights; Fig. 3D). This volatility in the sign of the parameter estimates for *K* indicates that any turning behaviour that is not already modelled by the PN element of the PN+PP simulations is not being consistently modelled by their PP element either, so is likely to represent noise. Hence, given that *K* > 0 steers flight towards the target, whereas *K* < 0 steers flight away from it, there is no evidence to indicate that a PP element supplements PN in commanding steering towards the target. Conversely, the observation that the parameter estimates for *N* were almost always positive in the successful PN simulations (median *N*: 1.2; 1^st^, 3^rd^ quartiles: 0.5, 1.4; sign test: *p* < 0.001; *n* = 18 flights; Fig. 3C), and were similar in the PN+PP simulations (median *N*: 1.1; 1^st^, 3^rd^ quartiles: 0.5, 1.5; sign test: *p* = 0.003; *n* = 20 flights; Fig. 3C), provides strong positive evidence that steering towards the target is indeed based on feeding back the line-of-sight rate *λ* rather than the deviation angle *δ*.

Several of the successful PN simulations had parameter estimates *N* ≪ 1, so predict little turning behaviour at all (Fig. 4). These simulations provide no positive evidence for feedback of *λ*, but all 4 simulations with values of *N* falling below the 1^st^ quartile correspond to nearly straight sections of flight (Fig. 4D,K,L,N), for which there is little turning behaviour to explain, and for which parameter estimation is therefore unreliable. Conversely, all 11 simulations with values of *N* falling between the 1^st^ and 3^rd^ quartiles were from flights involving a substantial amount of horizontal turning (Figs. 4C,E-I,M,O-R), which proportional feedback of *λ* successfully explains. To check whether PN guidance could also capture the altitudinal component of the Gyrfalcon flights, we re-fitted all of the PN simulations in 3D (Fig. 5). Although the number of flights that could be modelled successfully under PN dropped to 12/20 in 3D, comprising 734 m and 69.4 s of flight fitted at 1.2% error tolerance, the parameter estimates in the *n* = 12 successfully-fitted 3D simulations (median *N*: 1.0; 1^st^, 3^rd^ quartiles: 0.2, 1.4) were similar to those of the same flights in 2D (median *N*: 1.2; 1^st^, 3^rd^ quartiles: 0.6, 1.8), and were not significantly higher or lower in either case (sign test: *p* = 0.39). We therefore focus the remainder of our reporting on the 2D simulations.

**Figure 4.**
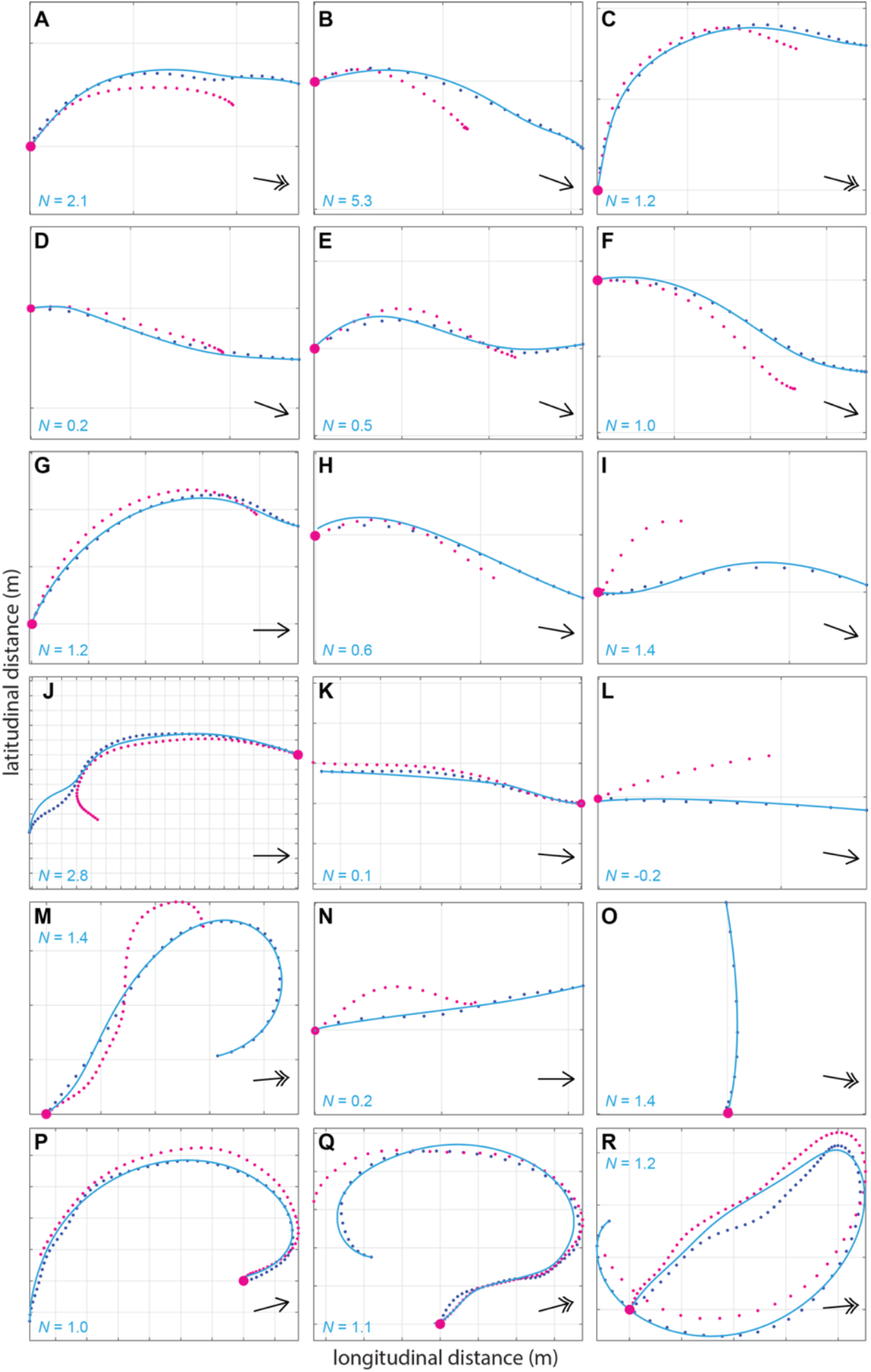
Two-dimensional (2D) attack trajectories for the 18/20 flights from *n* = 13 Gyrfalcons that were successfully modelled under proportional navigation (PN) guidance. Panels show the measured trajectories of the target (magenta points) and attacker (blue points), overlain with the longest simulation fitted to within 1.0% error tolerance (blue lines) in 2D. The corresponding parameter estimate for *N* is displayed on each plot. Note that among the 9 short dashes (panels A-I), 7 flights (panels A-G) are modelled in their entirety from target launch to intercept; the other 9 flights (panels J-R) each correspond to the terminal phase of an extended chase. Simulations with values of *N* falling beneath the 1^st^ quartile (*N* < 0.5) coincide with nearly straight sections of flight (panels D,K,L,N), for which parameter estimation is unreliable. Simulations with values of *N* falling between the 1^st^ and 3^rd^ quartiles (0.5 ≤ *N* ≤ 1.4) involve a substantial amount of turning that the model successfully explains (panels C,E-I,M,O-R). Black arrows display mean wind direction; double headed arrows correspond to wind speeds >20 km h^−1^; gridlines at 10m spacing. See Fig. S1 for the remaining 2/20 flights that were not successfully modelled under PN.

**Figure 5.**
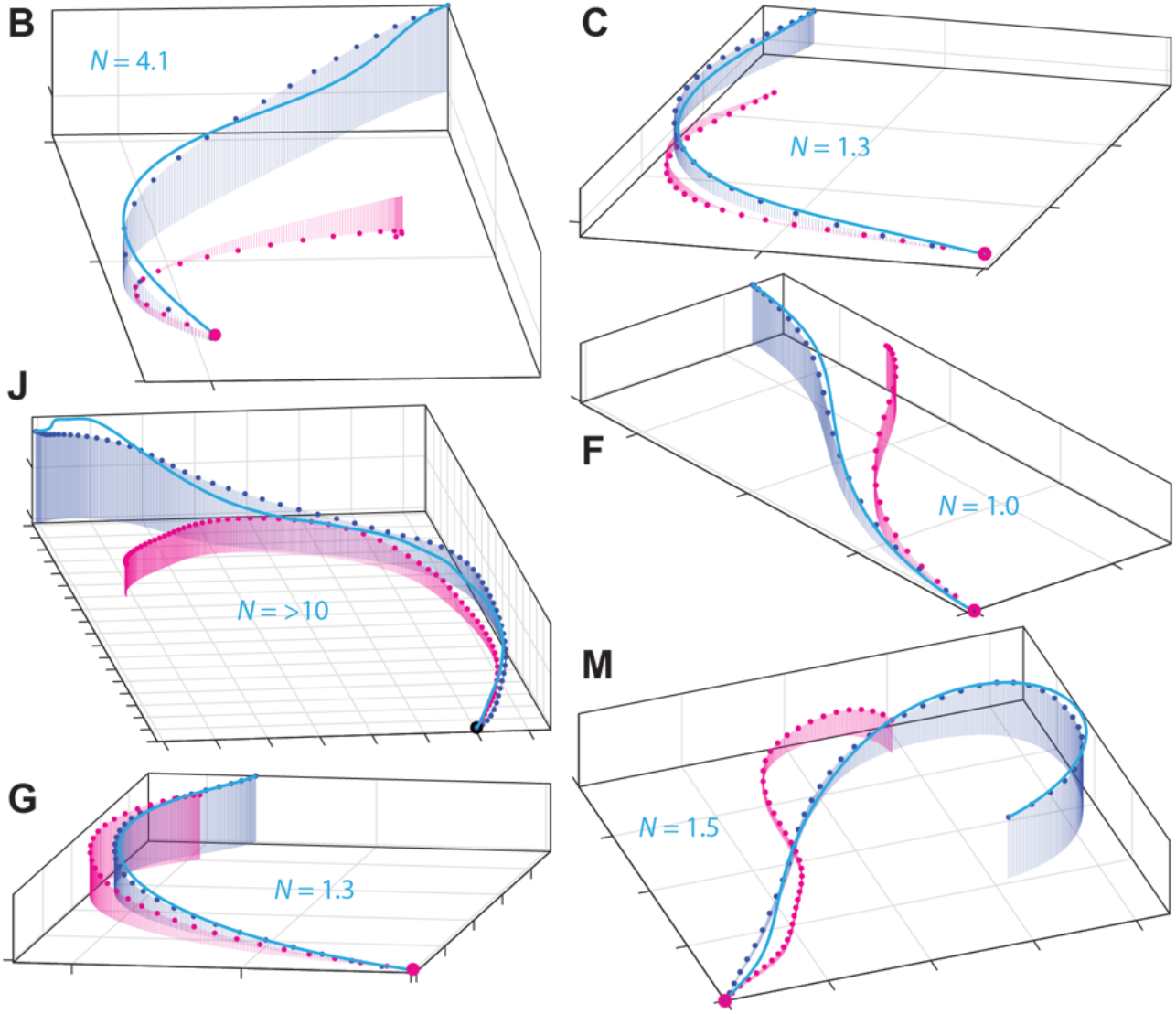
Three-dimensional (3D) attack trajectories for the subset of 6 Gyrfalcon flights involving the greatest altitudinal change among the 12/20 that were successfully modelled under proportional navigation (PN) guidance. Panels show the measured trajectories of the target (magenta points) and attacker (blue points), overlain with the longest simulation fitted to within 1.2% error tolerance (blue lines) in 3D. The corresponding parameter estimate for *N* is displayed on each plot. Panels B, C, F, G correspond to short dashes for which almost the entire flight was modelled from target launch to intercept. Panel letters match those in Fig. 4; gridlines at 10m spacing.

### PN models both short dashes and extended chases in their terminal phase

The 20 Gyrfalcon flights comprised 10 short dashes lasting from 3 to 7 s (Fig. 4A-I), and 10 extended chases lasting from 18 to 223 s (Fig. 4J-R). Typically, these short dashes correspond to the birds’ maiden flights (Table S1), whereas the extended chases correspond to their second flights (Table S1), which is because the pilot made less of an attempt to evade capture on the maiden flight. We found no evidence of any systematic increase or decrease in the parameter estimates for *N* between the Gyrfalcons’ maiden and second flights, for the small subsample of *n = 6* individuals for which paired data was available (sign test: *p* = 0.69). Of the 10 short dashes, 7 were modelled successfully from target launch to intercept (Fig. 4A-G), whilst 2 were modelled successfully for two-thirds of the total distance flown (Fig. 4H,I). For the 9/10 extended chases that were modelled successfully (Fig. 4J-R), the simulations only captured the terminal phase of the attacks (median duration fitted: 8.2 s; first, third quartiles: 3.1, 12.3 s). This nevertheless represents a substantial amount of flight fitted by distance (median distance fitted: 86.9 m; first, third quartiles: 19.9 m, 146.5 m), because of the high speeds reached by the end of a chase.

### Naïve Gyrfalcons operate at lower navigation constants than experienced Peregrines

All 13 flights recorded previously from experienced Peregrines (Brighton et al. 2017) were modelled successfully in 2D under PN or PN+PP (Fig. S2; Table S3), whereas only 11/13 of these flights were modelled successfully under PP (Table S3). Moreover, whereas the PN simulations fitted 1517 m and 99.6 s of flight at 1.0% error tolerance, the PP simulations fitted only 927 m and 57.8 s of flight, for the same number of estimated guidance parameters (Table S3). This confirms our earlier finding (Brighton et al. 2017) that PN is the better supported of the two pure guidance laws in Peregrines. In comparison, the simulations under PN+PP fitted 1740 m and 131.2 s of flight successfully at 1.0% error tolerance (Table S3). The addition of a PP term to the PN model for Peregrines therefore increased the duration of flight fitted by a factor of 1.3, but for a doubling in the number of estimated guidance parameters. It follows that the Peregrines’ attack trajectories are more economically modelled by PN than by PN+PP, which is the same conclusion as we have reached above for the Gyrfalcons.

The parameter estimates for *N* in the Gyrfalcons (median *N*: 1.2; 1^st^, 3^rd^ quartiles: 0.5, 1.4) were systematically lower (Fig. 6) than those in the Peregrines (median *N*: 2.8; 1^st^, 3^rd^ quartiles: 1.6, 3.1). This difference was statistically significant (Mood’s median test), both when treating repeated measures from the same individual as independent datapoints (*χ*^2^(1, *n* = 31) = 7.30; *p* = 0.007), and when analysing only the median values of *N* for each individual to eliminate pseudo-replication (*χ*^2^(1, *n* = 16) = 5.33; *p* = 0.02). The statistically significant difference in the distribution of *N* between the two species has a profound effect on the chase dynamics, as can be shown by simulating the intercept trajectories that the Gyrfalcons would have followed had they used the median value of *N* for the Peregrines, and *vice versa* (Fig. 7). It is clear by inspection that the Gyrfalcons would have intercepted the target sooner had they followed the trajectory commanded at *N* = 2.8 typical of Peregrines, and conversely that the Peregrines would not have intercepted the target as soon as they did had they followed the trajectory commanded at *N* = 1.2 typical of Gyrfalcons. This begs the question of why the Gyrfalcons operated at such low values of *N*, which we tackle from various perspectives in the Discussion, having first considered our key findings and their limitations.

**Figure 6.**
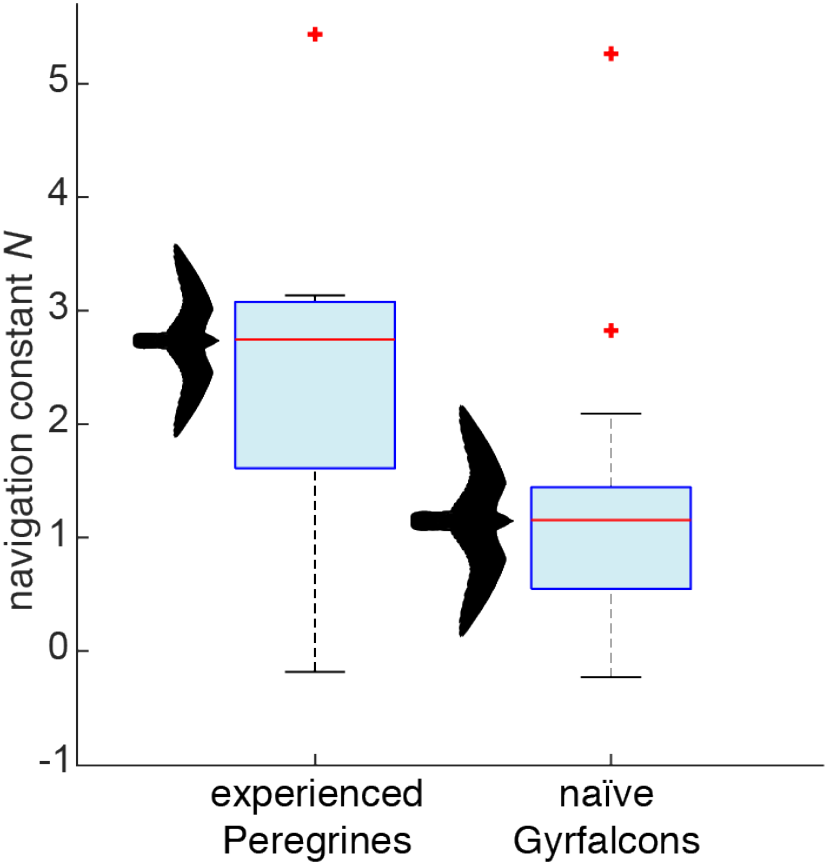
Box-and-whisker plots comparing parameter estimates for *N* in proportional navigation (PN) guidance models fitted independently to the 13 attack flights from *n* = 4 Peregrines, and 18 attack flights from *n* = 13 naïve Gyrfalcons; all in pursuit of manoeuvring targets. The centre line of each box denotes the median for all flights; the lower and upper bounds of the box denote the 1^st^ and 3^rd^ quartiles; crosses indicate outliers falling >1.5 times the interquartile range beyond the 1^st^ or 3^rd^ quartile (one extreme outlier for the Peregrines not shown); whiskers extend to the farthest datapoints not treated as outliers.

**Figure 7.**
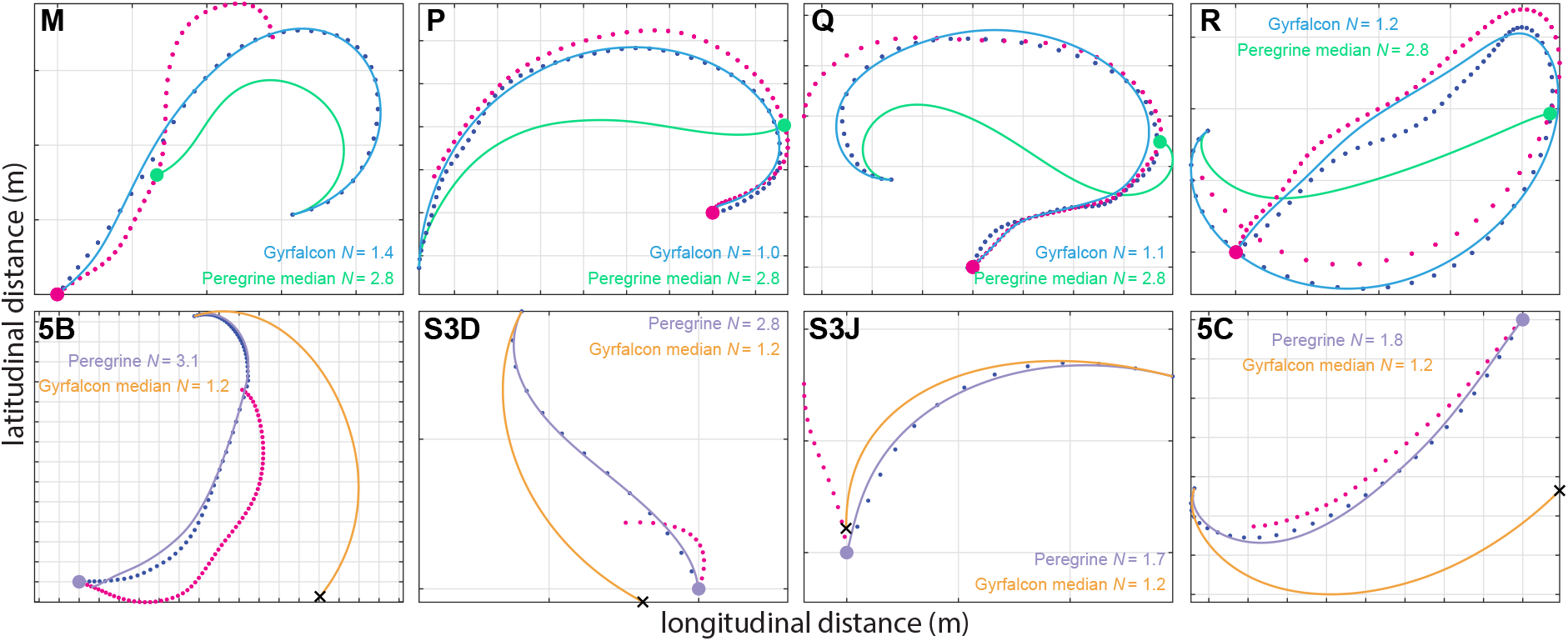
Effect of the navigation constant *N* on the dynamics of proportional navigation (PN) guidance. Upper row: selection of 4 successfully modelled Gyrfalcon flights involving substantial turning, showing the measured trajectory of the target (magenta dots) and attacker (blue dots) overlain with the best-fitting trajectory under PN guidance at the value of *N* displayed on the panel (blue line), and with the trajectory that would have been followed for the same initial conditions and target motion at the median value of *N* = 2.8 for Peregrines (green line). Green dot shows the predicted point of intercept had the Gyrfalcon used the median value of *N* for Peregrines; note that this is always sooner than the actual point of intercept (magenta dot). Lower row: selection of 4 successfully modelled Peregrine flights involving substantial turning, showing the measured trajectory of the target (magenta dots) and attacker (blue dots) overlain with the best-fitting trajectory under PN guidance at the value of *N* displayed on the panel (lilac line), and with the trajectory that would have been followed for the same initial conditions and target motion at the median value of *N* = 1.2 for Gyrfalcons (orange line). Lilac dot shows the actual point of intercept; black cross shows the predicted position of the bird had the Peregrine used the median value of *N* for Gyrfalcons; note that this is always at some significant distance from the target. Panel letters match those in Figs. 3, S2; gridlines at 10 m spacing.

## DISCUSSION

### Key scientific findings and their limitations

There are two key findings of this work: first, that naïve Gyrfalcons chase aerial targets as if using the same PN guidance law found previously in experienced Peregrines (Brighton et al. 2017); and second, that they do so at a lower value of the navigation constant *N* (Fig. 3). The first of these key findings is robustly confirmed by showing that PN models the data more successfully than PP and more economically than PN+PP. It is plausible that the data might be even better modelled by some alternative guidance law that we have not yet tested, but as there are only a limited number of variables that can be fed back to command steering in a particle model of interception, we think it likely that any such guidance law would be a variant of PN. The second of these key findings is robustly confirmed using non-parametric statistics that demonstrate a systematic difference in the navigation constant *N* between the Gyrfalcons (median *N*: 1.2; 1^st^, 3^rd^ quartiles: 0.5, 1.4) and the Peregrines (median *N*: 2.8; 1^st^, 3^rd^ quartiles: 1.6, 3.1) for identically-analysed data collected using identical GPS loggers. Its primary limitations are first that the experiments with Peregrines used a towed lure rather than a Roprey model, and second that the sample of Peregrines comprised only *n* = 4 individuals. However, as we have previously reported similar navigation constants (median: *N* = 2.6; 1^st^, 3^rd^ quartiles: 1.7, 3.3) in an independent sample of *n* = 3 experienced Peregrines attacking stationary targets (Brighton et al. 2017), we are confident that this result can be generalised.

### Technical limitations

It is important to emphasise that we have not identified a unique navigation constant *N* for either species. Rather, we have identified distinct intervals within which their respective values of *N* typically fall. Moreover, whilst we have reported a unique estimate of *N* for each flight, a different parameter estimate would have been reported had a different start point been chosen for the simulation. The selected start points are those which maximise the amount of flight fitted at the specified error tolerance, and therefore represent an objective compromise between goodness of fit and amount of data modelled. It is uncertain how much of the variability in our parameter estimates for *N* is the result of measurement error as opposed to genuine behavioural flexibility. In particular, the use of GPS loggers recording speed and position at 5 Hz precludes perfect synchronization of the attacker and target trajectories and hence estimation of the sensorimotor delay, so it is plausible that data collected at higher spatiotemporal precision would show lower variability in the parameter estimates for *N*. Finally, our modelling only accounts for the influence of wind only insofar as the simulated groundspeed is matched to that of the measured groundspeed. The simulations therefore treat any distortions of the bird’s track due to the wind as if these were produced by the attacker’s own steering commands, although the effect of this is mitigated in close pursuit by the fact that the attacker and its target are subject to the same wind. Nevertheless, gusting winds might perhaps explain some of the more prominent inflections in the measured flight trajectories that are not captured by simulations fitted over longer distances (e.g. Fig. 4J) or at higher wind speeds (e.g. Fig. 4R).

### Physical and physiological constraints

For a given line-of-sight rate, the higher values of *N* in Peregrines are expected to command turning at approximately twice the angular rate of the lower values of *N* in Gyrfalcons. Minimum turn radius scales linearly with wing loading, *W* (Taylor and Thomas 2014), which is ~6.0 kg m^−2^ in Gyrfalcons compared to ~5.3 kg m^−2^ in Peregrines (Pennycuick et al. 1994). Gyrfalcons are therefore expected to be less manoeuvrable than Peregrines, with a ~13% larger minimum turn radius. They are also expected to be less agile, because flight speed scales as *W^1/2^* (Taylor and Thomas 2014), such that maximum turn rate scales as *W^−1/2^*, and is therefore expected to be ~6% lower in Gyrfalcons than Peregrines. In principle, this physical constraint could force the navigation constant *N* to be lower in Gyrfalcons, but the expected difference in their agility is too small to explain the twofold difference in *N* that we observe. Furthermore, there is no evidence that physical constraints on turning influenced the shape of the recorded trajectories. Rather than turning as tightly as possible on a circular arc, the birds instead followed a curved trajectory of increasing or decreasing radius, characteristic of the time history of turning commanded under PN guidance at higher or lower values of *N*, respectively (Fig. 4). The observed flight trajectories are similarly inconsistent with an old hypothesis proposed by Tucker (Tucker 2000), who argued that the curved attack trajectories of falcons could be generated by steering so as to hold the target’s image on the laterally directed central fovea of the left or right eye whilst holding the head in line with the body (Tucker et al. 2000). This constraint is expected to produce a trajectory in which the deviation angle *δ* between the attacker’s velocity and its line-of-sight to target is held constant at approximately ± 40°, but there is no evidence of this from the data (see Fig. 2; see also (Kane and Zamani 2014).

### A functional account of the observed variation in navigation constant

In principle, the variation in *N* within and between species (Fig. 6) might be explained as a behavioural response under linear-quadratic optimal guidance theory (Shneydor 1998; Siouris 2004), which predicts an optimum value of *N* = 3 *v_c_*/(*v* cos *δ*) for attacks on non-manoeuvring targets minimizing overall steering effort. In practice, the median values of the ratio *v_c_*/(*v* cos *δ*) for each flight did not differ significantly between the two species (*χ*^2^(1, *n* = 31) = 1.55; *p* = 0.21), and nor were they significantly related to *N* in a bisquare robust regression controlling for species (*t*(28) = 0.46; *p* = 0.69). We therefore find no evidence that the statistically significant difference in the parameter estimates for *N* between the Peregrines and Gyrfalcons reflects a direct functional response to variation in the rate at which they closed range on the target relative to their own approach speed. This conclusion is based on classical theory for non-manoeuvring targets, and therefore takes no account of the target’s manoeuvres. Nevertheless, the tortuosity of the lure’s path, defined as its overall path length divided by the straight line distance from start to finish, was similar in the experiments with Peregrines (median tortuosity: 1.6; 1^st^, 3^rd^ quartiles: 1.3, 2.4) and Gyrfalcons (median tortuosity: 1.7; 1^st^, 3^rd^ quartiles: 1.3, 2.5), so we conclude that differences in target manoeuvres are unlikely to explain the variation in *N* that we observed between species. Even so, an identical target manoeuvre will produce a greater angular change in the line-of-sight vector the closer the attacker is to its target, thereby demanding a higher rate of turning under PN. This is important, because whereas the Gyrfalcons took off together with the target at a range of ~20 m, the Peregrines were launched separately and sometimes at a range of >100 m. When operating at close range, PN guidance is less prone to being thrown off by erratic target manoeuvres if *N* is low (Brighton and Taylor 2019), so it is plausible that the lower values of *N* found in the Gyrfalcons might reflect a behavioural response to their proximity to the target at the initiation of the attack.

### An adaptive account of the observed variation in navigation constant

A complementary adaptive argument can also be made at species level. Whereas Peregrines have a very flexible diet (Cade 1982), Gyrfalcons depend heavily on a single prey type, with Ptarmigan (*Lagopus* spp.) comprising 74% of the catch on average across 17 studies recording 66,726 individual prey items over most of the Gyrfalcon’s range (Nielsen and Cade 2017). Ptarmigan have a high wing loading of ~9.7 kg m^−2^ (Greenewalt 1962), which is ~62% higher than that of a Gyrfalcon. They are therefore intrinsically fast fliers, with large flight muscles that provide rapid acceleration in an explosive take-off, and short wings that make them less well adapted to sustained aerobic flight (Pennycuick et al. 1994). The prolonged tail-chasing behaviour that is typical of Gyrfalcons (Cade 1982) is therefore thought to serve to tire their prey and allow capture (Pennycuick et al. 1994). Such behaviour is promoted by the low values of *N* found in Gyrfalcons (Fig. 7), because PN with a low navigation constant *N* ≈ 1 commands turning at an angular rate *γ* approximately equal to the line-of-sight rate *λ*. Once a tail chase has been initiated, this will tend to keep the attacker flying behind its target, and will not tend to be thrown off too far by the erratic jinking manoeuvres that are characteristic of the evasive flight of Ptarmigan and many other prey (Mills et al. 2018; Mills et al. 2019). The low values of *N* found in Gyrfalcons therefore make sense from an adaptationist perspective if the function of their PN guidance is to pursue the prey doggedly until it tires. This is in contrast to the higher values of *N* found in Peregrines, which make sense if the function of their PN guidance is to exploit the speed and manoeuvrability acquired through stooping in order to intercept prey quickly and efficiently (Mills et al. 2018; Mills et al. 2019).

### Evolutionary and ontogenetic implications

We found no evidence that the value of *N* changed systematically between the Gyrfalcons’ first and second flights, but it is reasonable to assume that they would have learned to tune their guidance over longer timescales. For example, although wild Gyrfalcons do not usually stoop, falconers commonly train captive birds to do so (Tucker et al. 1998), which implies a degree of flexibility in their guidance. Likewise, whereas naïve Gyrfalcons tend to fly directly at their prey, experienced birds seem to anticipate their prey’s behaviour. Even so, our finding that the maiden attack flights of naïve Gyrfalcons are well modelled under PN guidance strongly suggests that this behavioural algorithm is embedded in a hardwired guidance loop. Moreover, the fact that the same form of guidance law models the attack flights of experienced Peregrines is consistent with the hypothesis that PN guidance is ancestral to the clade comprising the Peregrines and Hierofalcons, of which the Gyrfalcon is a member (Wink 2018). Formal confirmation of this hypothesis would require the same behavioural algorithm to be identified in another Hierofalcon, or in a close outgroup such as the Merlin or Hobby falcons, both of which specialise in aerial pursuit. Because PN guidance commands turning in proportion to the line-of-sight rate of the target, which is defined in an inertial frame of reference, we would expect any such birds using PN to share a common sensorimotor architecture that fuses sensory input from the visual and vestibular systems to obtain the line-of-sight rate, and that uses the resulting signal to generate motor commands to the flight muscles controlling the wings and tail. Alternatively, it is possible that the distant landscape might serve as a visual proxy for the inertial frame of reference in aerial hunters, with target motion being measured directly with respect to the visual background; see also (Kane and Zamani 2014). It would therefore be of particular interest to test whether PN guidance also models the attack behaviours of the more distantly related Kestrels, given their specialism on terrestrial prey with no distant visual background against which to assess prey motion.

## Acknowledgements

We thank Jimmy Robinson and Tom Spink for falconry work, plus their interns, Adam Vacek, Sander Gielen, and Daniel Clark. We thank Remy Van Wijk and Matt Aggett for piloting the Roprey. We thank Barbro Fox for her hospitality, and Sofía Miñano Gonzalez for valuable comments on the manuscript.

## Author contributions

CB, GT and NF conceived the study. All authors contributed to field experiments. CB analysed the GPS data. CB and GT performed statistical analysis and wrote the paper. All authors commented on and approved the final version of the manuscript.

## References

Bengtson SA. 1971. Hunting methods and choice of prey of gyrfalcons falco-rusticolus at myvatn in northeast iceland. Ibis. 113(4):468–&.

Brighton CH, Taylor GK. 2019. Hawks steer attacks using a guidance system tuned for close pursuit of erratically manoeuvring targets. Nat Commun. 10.

Brighton CH, Thomas ALR, Taylor GK. 2017. Terminal attack trajectories of peregrine falcons are described by the proportional navigation guidance law of missiles. P Natl Acad Sci USA. 114(51):13495–13500.

Cade TJ. 1982. The falcons of the world. Ithaca, New York, USA.: Cornell University Press.

Cresswell W. 1996. Surprise as a winter hunting strategy in sparrowhawks accipiter nisus, peregrines falco peregrinus and merlins f-columbarius. Ibis. 138(4):684–692.

Garber CS, Mutch BD, Platt S. 1993. Observations of wintering gyrfalcons (falco-rusticolus) hunting sage grouse (centrocercus-urophasianus) in wyoming and montana USA. J Raptor Res. 27(3):169–171.

Greenewalt CH. 1962. Dimensional relationships for flying animals. Smithsonian Miscellaneous Collections Smithsonian Institution. 144.

Hein AM, Altshuler DL, Cade DE, Liao JC, Martin BT, Taylor GK. 2020. An algorithmic approach to natural behavior. Curr Biol. 30(11):R663–R675.

Jarvis ED, Mirarab S, Aberer AJ, Li B, Houde P, Li C, Ho SYW, Faircloth BC, Nabholz B, Howard JT et al. 2014. Whole-genome analyses resolve early branches in the tree of life of modern birds. Science. 346(6215):1320–1331.

Kane SA, Zamani M. 2014. Falcons pursue prey using visual motion cues: New perspectives from animal-borne cameras. J Exp Biol. 217(2):225–234.

Mills R, Hildenbrandt H, Taylor GK, Hemelrijk CK. 2018. Physics-based simulations of aerial attacks by peregrine falcons reveal that stooping at high speed maximizes catch success against agile prey. Plos Comput Biol. 14(4):e1006044.

Mills R, Taylor GK, Hemelrijk CK. 2019. Sexual size dimorphism, prey morphology and catch success in relation to flight mechanics in the peregrine falcon: A simulation study. J Avian Biol. 50(3).

Nielsen OK, Cade TJ. 2017. Gyrfalcon and ptarmigan predator-prey relationship. In: Anderson DL, McClure CJW, Franke A, editors. Applied raptor ecology: Essentials from gyrfalcon research. Boise, Idaho, USA: The Peregrine Fund.

Pennycuick CJ, Fuller MR, Oar JJ, Kirkpatrick SJ. 1994. Falcon versus grouse - flight adaptations of a predator and its prey. J Avian Biol. 25(1):39–49.

Potapov E, Sale R. 2005. The gyrfalcon. Yale University Press.

Prum RO, Berv JS, Dornburg A, Field DJ, Townsend JP, Lemmon EM, Lemmon AR. 2015. A comprehensive phylogeny of birds (aves) using targeted next-generation DNA sequencing. Nature. 526(7574):569–U247.

Shneydor NA. 1998. Missile guidance and pursuit: Kinematics, dynamics and control. Woodhead Publishing Limited.

Siouris GM. 2004. Missile guidance and control systems. Springer-Verlag.

Taylor GK, Thomas ALR. 2014. Evolutionary biomechanics: Selection, phylogeny, and constraint. Ox Ecol Ev.1-152.

Tucker VA. 2000. The deep fovea, sideways vision and spiral flight paths in raptors. J Exp Biol. 203(24):3745–3754.

Tucker VA, Cade TJ, Tucker AE. 1998. Diving speeds and angles of a gyrfalcon (falco rusticolus). J Exp Biol. 201(13):2061–2070.

Tucker VA, Tucker AE, Akers K, Enderson HJ. 2000. Curved flight paths and sideways vision in peregrine falcons (falco peregrinus). J Exp Biol. 203(24):3755–3763.

White CM, and Nelson, R. W. 1991. Hunting range and strategies of tundra breeding peregrine and gyrfalcons observed from a helicopter. J Raptor Res. 25:49–62.

White CM, Weeden RB. 1966. Hunting methods of gyrfalcons and behavior of their prey (ptarmigan). Condor. 68(5):517–&.

Wink M. 2018. Phylogeny of falconidae and phylogeography of peregrine falcons. Ornis Hungarica. 26(2):27–37.

Woodin N. 1967. Observations on gyrfalcons (falco rusticolus) breeding near lake myvatn, iceland. Raptor Research. 14(4):97–124.

