## Supplementary figures S1, S2 and tables S1-S3 for "Attack behaviour in naïve Gyrfalcons is modelled by the same guidance law as in Peregrines, but at a lower guidance gain"

### Supplementary Information

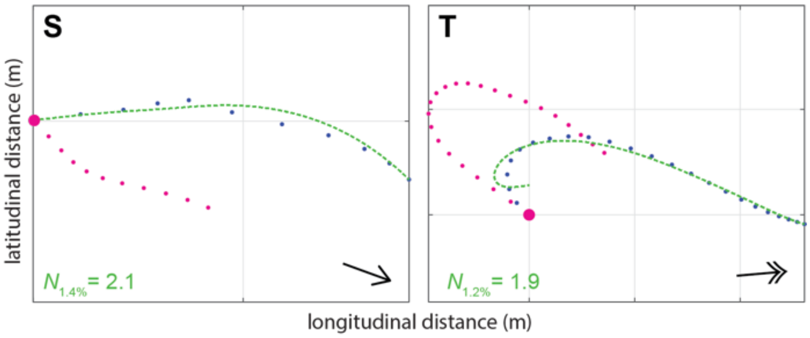


**Figure S1**. Two-dimensional (2D) attack trajectories for the 2/20 flights from $n = 13$ Gyrfalcons that were not successfully modelled under proportional navigation (PN) guidance. Panels show the measured trajectories of target (magenta points) and attacker (blue points), overlain with the simulation with the lowest relative error lasting ≥ 2 s (green dashed line). The corresponding parameter estimate for $N$ is displayed on each plot, with a subscript indicating the relative error achieved. Black arrows display mean wind direction; double headed arrows correspond to wind speeds >20 km $h^{-1}$; gridlines at 10m spacing.

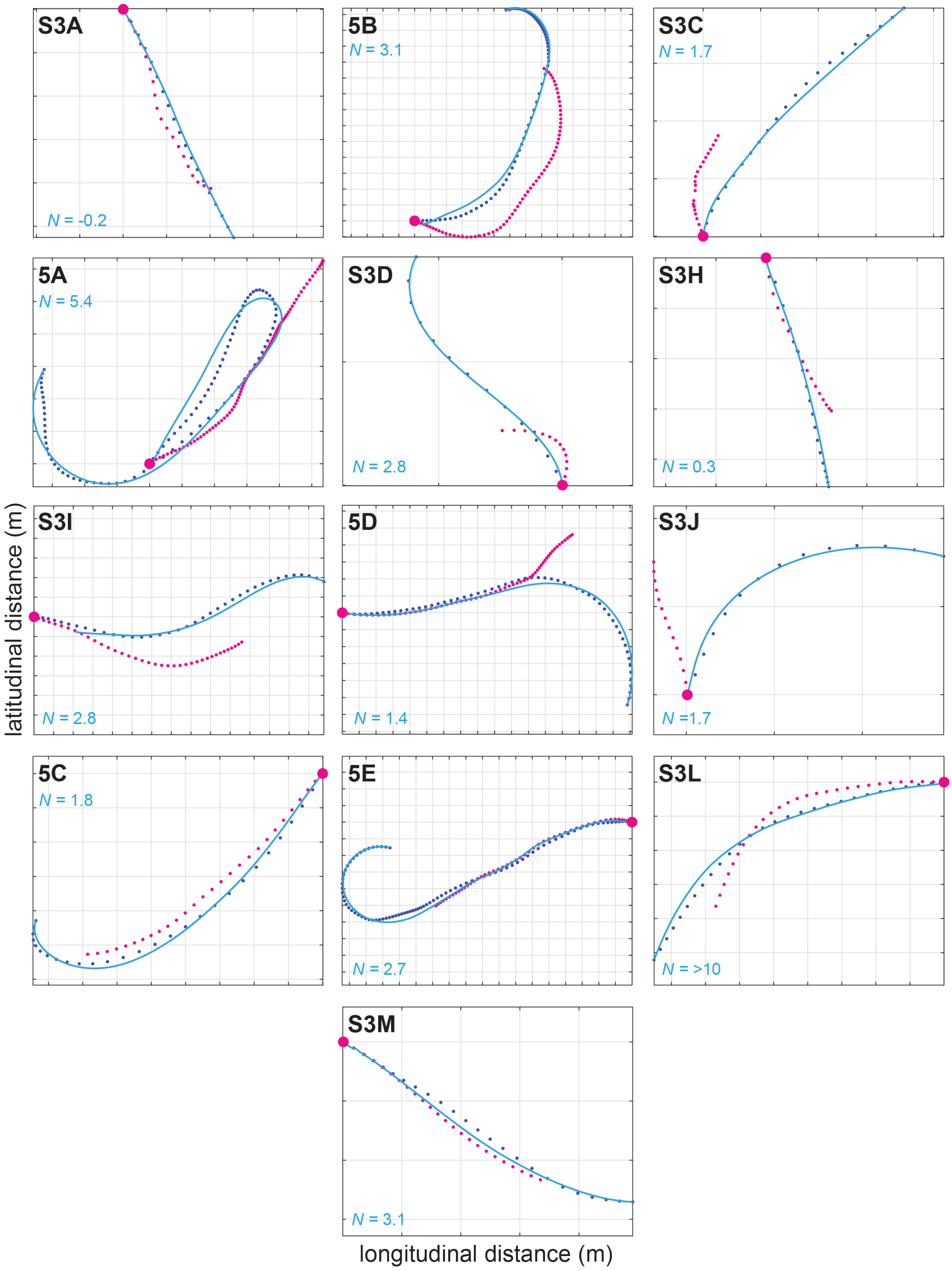


**Figure S2**. Two-dimensional (2D) attack trajectories for the 13/13 flights from $n = 4$ Peregrines that were successfully modelled under proportional navigation (PN) guidance. Panels show the measured trajectories of the target (magenta points) and attacker (blue points), overlain with the longest simulation fitted to within 1.0% error tolerance (blue lines) in 2D. The corresponding parameter estimate for $N$ is displayed on each plot. Note that the only simulations with values of $N$ falling beneath the 3[rd] quartile for the Gyrfalcons ($N < 1.4$) coincide with nearly straight sections of flight (panels S3A, S3H), for which parameter estimation is unreliable. Panel labels correspond to the numbering and lettering of the figure panels in the original paper from which the measured trajectory data are drawn. Gridlines at 10m spacing.

### Tables

| date | bird ID | bird flight | flight | type | pedigree | sex | mass (kg) |
|---|---|---|---|---|---|---|---|
| 10/08/17 | 10 | 2nd | S | chase | Gyrfalcon | ♀ | 1.295 |
| 10/08/17 | 8 | 2nd | J | chase | Gyrfalcon | ♂ | 0.939 |
| 25/08/17 | 11 | 1st | K | chase | Gyrfalcon | ♀ | 1.098 |
| 25/08/17 | 13 | 1st | A | dash | Gyrfalcon | ♂ | 0.850 |
| 25/08/17 | 16 | 1st | H | dash | Gyrfalcon | ♀ | 1.179 |
| 25/08/17 | 18 | 1st | B | dash | Gyrfalcon | ♀ | 1.144 |
| 25/08/17 | 17 | 1st | L | chase | Gyrfalcon | ♂ | 0.944 |
| 25/08/17 | 21 | 1st | C | dash | Gyrfalcon | ♂ | 0.871 |
| 25/08/17 | 20 | 1st | D | dash | Gyrfalcon | ♀ | 1.164 |
| 25/08/17 | 22 | 1st | E | dash | Gyrfalcon | ♂ | 0.877 |
| 25/08/17 | 23 | 1st | F | dash | Gyrfalcon | ♂ | 0.858 |
| 25/08/17 | 73 | 1st | I | dash | 7/8th Gyr-Saker | ♂ | 0.856 |
| 25/08/17 | M4 | 1st | G | dash | 7/8th Gyr-Saker | ♀ | 1.136 |
| 26/08/17 | 13 | 2nd | T | dash | Gyrfalcon | ♂ | 0.838 |
| 26/08/17 | 18 | 2nd | M | chase | Gyrfalcon | ♀ | 1.143 |
| 26/08/17 | 17 | 2nd | N | chase | Gyrfalcon | ♂ | 0.937 |
| 26/08/17 | 23 | 2nd | O | chase | Gyrfalcon | ♂ | 0.873 |
| 26/08/17 | 73 | 2nd | P | chase | 7/8th Gyr-Saker | ♂ | 0.872 |
| 26/08/17 | M4 | 2nd | Q | chase | 7/8th Gyr-Saker | ♀ | 1.129 |
| 26/08/17 | 13 | 3rd | R | chase | 7/8th Gyr-Saker | ♂ | 0.838 |

**Table S1.** Details of the $n = 13$ Gyrfalcons used in the 20 flights meeting quality control for accuracy of the GPS data. The letter code for each flight corresponds to the lettering of all applicable figure panels.

| date | bird | flight | PP | | | | PN | | | | PN+PP | | | | |
|---|---|---|---|---|---|---|---|---|---|---|---|---|---|---|---|
| | | | $K$ (s$^{-1}$) | duration (s) | distance (m) | relative error (%) | $N$ | duration (s) | distance (m) | relative error (%) | $N$ | $K$ (s$^{-1}$) | duration (s) | distance (m) | relative error (%) |
| 10/08/17 | 14 | S | 0.9 | 2.0 | 18.8 | 0.70 | n.s. | n.s. | n.s. | n.s. | -0.3 | 1.1 | 5.4 | 56.5 | 0.74 |
| 10/08/17 | 1 | J | 0.6 | 7.8 | 171.3 | 0.88 | 2.8 | 10.6 | 212.7 | 1.04 | 1.0 | 0.3 | 13.6 | 225.3 | 1.04 |
| 25/08/17 | 2 | K | 0.1 | 10.2 | 107.4 | 0.99 | 0.1 | 6.4 | 65.7 | 0.81 | -0.1 | 0.1 | 10.2 | 107.4 | 0.65 |
| 25/08/17 | 4 | A | 1.1 | 3.2 | 18.8 | 1.01 | 2.1 | 5.4 | 28.9 | 0.86 | 1.0 | 0.2 | 5.4 | 28.9 | 0.56 |
| 25/08/17 | 5 | H | 1.0 | 4 | 20.9 | 1.04 | 0.6 | 2.6 | 17.1 | 1.03 | 1.0 | 0.9 | 4 | 20.9 | 0.28 |
| 25/08/17 | 6 | B | n.s. | n.s. | n.s. | n.s. | 5.3 | 4.2 | 22.4 | 0.68 | 1.5 | 0.4 | 4.2 | 22.4 | 0.23 |
| 25/08/17 | 7 | L | -0.1 | 2.2 | 14.8 | 0.43 | -0.2 | 2.2 | 14.8 | 0.52 | 1.4 | -1.0 | 3.4 | 24.6 | 0.92 |
| 25/08/17 | 8 | C | 8.8 | 3.8 | 35.2 | 1.01 | 1.2 | 5 | 40.6 | 0.60 | 1.6 | -0.8 | 5.2 | 41.1 | 0.59 |
| 25/08/17 | 9 | D | 0.2 | 3.4 | 27.0 | 0.81 | 0.2 | 3.6 | 27.6 | 0.71 | 0.4 | -0.2 | 3.6 | 27.6 | 0.52 |
| 25/08/17 | 10 | E | n.s. | n.s. | n.s. | n.s. | 0.5 | 3.8 | 32.2 | 0.96 | 0.7 | -0.4 | 3.8 | 32.2 | 0.78 |
| 25/08/17 | 11 | F | n.s. | n.s. | n.s. | n.s. | 1.0 | 4.2 | 37.9 | 0.61 | 1.5 | 0.3 | 4.2 | 37.9 | 0.51 |
| 25/08/17 | 12 | I | -0.3 | 2.4 | 8.4 | 0.70 | 1.4 | 3.2 | 14.3 | 1.04 | 1.8 | 1.75 | 4.0 | 18.1 | 0.28 |
| 25/08/17 | 13 | G | n.s. | n.s. | n.s. | n.s. | 1.2 | 6.4 | 57.3 | 0.85 | 2.2 | 2.5 | 7 | 59.2 | 0.65 |
| 26/08/17 | 4 | T | n.s. | n.s. | n.s. | n.s. | n.s. | n.s. | n.s. | n.s. | 1.6 | -1.8 | 3.6 | 31.3 | 0.45 |
| 26/08/17 | 6 | M | 1.2 | 6 | 63.2 | 1.04 | 1.4 | 8.2 | 86.9 | 0.35 | 1.8 | -0.2 | 8.4 | 89.8 | 0.53 |
| 26/08/17 | 7 | N | 0.2 | 3.2 | 21.6 | 0.68 | 0.2 | 3.2 | 21.6 | 0.81 | 0.0 | 0.2 | 3.2 | 21.6 | 0.67 |
| 26/08/17 | 9 | O | 1.9 | 2.6 | 10.0 | 0.64 | 1.4 | 2.6 | 10.0 | 0.41 | 1.2 | 0.22 | 2.6 | 10.0 | 0.42 |
| 26/08/17 | 12 | P | 1.7 | 13.6 | 162.6 | 1.00 | 1.0 | 12 | 148.3 | 0.51 | 1.0 | 0.2 | 18.8 | 202.4 | 1.01 |
| 26/08/17 | 13 | Q | 2.3 | 13 | 144.0 | 0.28 | 1.1 | 13.2 | 145.9 | 1.01 | -11.9 | 28.6 | 14.6 | 157.5 | 0.68 |
| 26/08/17 | 4 | R | n.s. | n.s. | n.s. | n.s. | 1.2 | 14.2 | 143.2 | 0.85 | 1.3 | -0.2 | 14.2 | 143.2 | 0.68 |

**Table S2.** Results of the two-dimensional (2D) guidance model fitting for the sample of 20 flights from $n = 13$ Gyrfalcons, under proportional pursuit (PP), proportional navigation (PN), and mixed PN+PP guidance. Flights marked "n.s." are those which could not be fitted to within the specified 1.0% error tolerance for $\geq 2$ s. The letter code for each flight corresponds to the lettering of all applicable figure panels. See text for details and definitions.

| date | bird | flight | PP | | | | PN | | | | PN+PP | | | | |
|---|---|---|---|---|---|---|---|---|---|---|---|---|---|---|---|
| | | | $K$ (s$^{-1}$) | duration (s) | distance (m) | relative error (%) | $N$ | duration (s) | distance (m) | relative error (%) | $N$ | $K$ (s$^{-1}$) | duration (s) | distance (m) | relative error (%) |
| 09/05/14 | Le | S3A | -0.2 | 3.6 | 58.5 | 0.51 | -0.2 | 3.6 | 58.5 | 0.44 | 0.3 | 0.7 | 8.8 | 121.6 | 1.00 |
| 28/04/14 | Le | 5B | -0.5 | 3.0 | 47.7 | 0.68 | 3.1 | 13.2 | 195.7 | 1.01 | 2.2 | 0.3 | 14.8 | 209.0 | 0.63 |
| 22/08/16 | Rn | S3C | -0.6 | 2.8 | 35.0 | 0.46 | 1.7 | 4.2 | 53.7 | 0.99 | 1.9 | 0.6 | 7.0 | 88.0 | 0.90 |
| 22/08/16 | Mo | 5A | -0.3 | 5.0 | 55.7 | 0.90 | 5.4 | 18.4 | 199.5 | 1.01 | 2.9 | 0.8 | 6.8 | 67.1 | 0.66 |
| 23/08/16 | Ra | S3D | n.s. | n.s. | n.s. | n.s. | 2.8 | 2.6 | 23.3 | 0.44 | 8.0 | -5.6 | 4.6 | 46.4 | 0.24 |
| 08/09/16 | Rn | S3H | 0.4 | 3.4 | 47.6 | 0.55 | 0.3 | 3.2 | 47.2 | 0.43 | 1.6 | 3.5 | 3.6 | 47.9 | 0.82 |
| 09/09/16 | Rn | S3I | 0.0 | 2.2 | 41.7 | 0.87 | 2.8 | 7.4 | 156.7 | 0.77 | 1.3 | 0.2 | 7.8 | 165.0 | 0.90 |
| 09/09/16 | Ra | 5D | 0.9 | 12.2 | 235.4 | 0.86 | 1.4 | 11.4 | 223.5 | 0.92 | 0.7 | 0.5 | 14.2 | 251.5 | 0.30 |
| 12/09/16 | Rn | S3J | n.s. | n.s. | n.s. | n.s. | 1.7 | 2.6 | 37.7 | 0.87 | 0.6 | 0.2 | 20.0 | 89.5 | 0.86 |
| 14/09/16 | Ra | 5C | 1.3 | 5.2 | 113.7 | 0.68 | 1.8 | 5.4 | 115.7 | 0.87 | 0.8 | 0.3 | 12.2 | 201.4 | 1.00 |
| 15/09/16 | Rn | 5E | 0.8 | 14.2 | 195.8 | 0.96 | 2.7 | 17.6 | 242.2 | 0.62 | 3.4 | -0.1 | 17.6 | 242.2 | 0.52 |
| 15/09/16 | Ra | S3L | -0.2 | 2.6 | 51.1 | 0.06 | >10 | 5.4 | 105.8 | 1.00 | 1.0 | 0.6 | 8.8 | 149.1 | 1.01 |
| 15/09/16 | Mo | S3M | -0.1 | 3.6 | 44.9 | 0.77 | 3.1 | 4.6 | 56.9 | 0.90 | 2.6 | -2.2 | 5.0 | 61.1 | 0.54 |

**Table S3.** Results of the two-dimensional (2D) guidance model fitting for the sample of 13 flights from $n = 4$ Peregrines, under proportional pursuit (PP), proportional navigation (PN), and mixed PN+PP guidance. Flights marked "n.s." are those which could not be fitted to within the specified 1.0% error tolerance for ≥ 2 s. The letter code for each flight corresponds to the lettering of all applicable figure panels. See text for details and definitions.
